# TAp63-regulated oncogenic long non-coding RNA-8 (*TROLL-8*) regulates human breast cancer progression through CPT1A-mediated fatty acid oxidation

**DOI:** 10.64898/2026.06.11.731679

**Authors:** Avani A. Deshpande, Marco Napoli, John H. Lockhart, Xiaobo Li, Min Liu, Lancia N. F. Darville, Bin Fang, John M. Koomen, Elsa R. Flores

## Abstract

Metabolic reprogramming is a crucial hallmark of cancer, supporting tumor growth and adaptation to cellular stress. Although fatty acid oxidation (FAO) has emerged as an important regulator in cancer, the mechanisms that control the FAO machinery remain poorly understood. Here, we demonstrate that the TAp63-regulated long non-coding RNA *TROLL-8* is a key regulator of FAO in breast cancer. Using metabolomics and protein microarray assays followed by immunoprecipitation-mass spectrometry, we mechanistically demonstrate that *TROLL-8* binds the FAO enzyme CPT1A and promotes the formation of a complex with ACSL1 and VDAC1, thereby enabling efficient fatty acid processing. Loss of *TROLL-8* destabilizes this complex, leading to impaired FAO, decreased metabolic fitness, and suppression of tumorigenic phenotypes, such as anchorage-independent growth. Notably, higher levels of CPT1A and VDAC1 are associated with worse survival in breast cancer patients. Given the emerging role of CPT1A in therapy resistance, these findings suggest that targeting *TROLL-8* could be a promising approach to selectively disrupt the hyperactive FAO machinery in breast cancers and other tumor types that rely on FAO for their progression.

## INTRODUCTION

Breast cancer remains the most frequently diagnosed tumor type and continues to be one of the leading causes of cancer-related death in women.^1^ Despite numerous advances in screening and therapies over the past few decades, disease progression and therapy resistance still pose significant challenges.^2^ One crucial biological process that has long been recognized as a hallmark of cancer progression is metabolic rewiring and reprogramming of tumor cells, which enable them to adapt to varying and often scarce nutrient availability and survive under chronic stress.^3,4^ Specifically, the rewiring of lipid metabolism has emerged as an essential driver of breast cancer progression.^5,6^ Indeed, in contrast to physiological conditions, where there is a finely regulated equilibrium between the *de novo* fatty acid synthesis (FAS) and the catabolism of fatty acid via β-oxidation,^5^ the increased rate of fatty acid oxidation (FAO) in cancer cells has been proven to support multiple pro-oncogenic features, including tumor growth, angiogenesis, immune evasion, and metastasis formation.^6,7^ Therefore, understanding the molecular mechanisms controlling FAO can provide novel therapeutic opportunities to hinder tumor progression reliant on FAO and ultimately improve the prognosis of these patients.

The p53 family of transcription factors suppresses tumor formation and progression by regulating several hallmarks of cancer, including cellular metabolism.^8,9^ In particular, we previously reported that the p63 isoform and p53 family member, TAp63, acts as an essential tumor and metastasis suppressor that controls breast cancer progression and metastasis and lipid metabolism.^10,11^ In fact, deletion of *TAp63* in mice leads to the spontaneous development of highly aggressive and metastatic tumors, with the most frequent being mammary adenocarcinomas with distant metastases to the lungs, liver, and brain.^10^ We found that the suppression of metastasis by TAp63 occurs primarily due to the TAp63-dependent transcriptional regulation of *Dicer*, thereby impairing the microRNA biogenesis pathway.^10^ In addition to tumor development, *TAp63^-/-^* mice also become obese and display other metabolic phenotypes, including liver steatosis, which are caused by the direct transcriptional regulation of *Sirt1*, *AMPKα*, and *LKB1* by TAp63,^11^ thus highlighting the crucial role of TAp63 at the intersection of metabolic reprogramming, lipid metabolism, and the development of metastatic breast cancer. To better understand the suppression of breast cancer by TAp63 in human patients, we previously performed^12^ a cross-species RNA-seq analysis comparing the *TAp63^-/-^* mammary adenocarcinoma mouse model with the human MCF10A breast cancer progression model,^13,14^ which led us to identify a group of 9 conserved TAp63-regulated long non-coding RNAs, or TROLLs.^12^ One of these lncRNAs, *TROLL-8*, exhibited increased RNA expression in both the mouse and human models of aggressive breast cancers, and we previously showed that downregulation of *TROLL-8* by siRNA reduced migration and invasion and increased apoptosis in breast cancer cells.^12^

Given these exciting results, we investigated the mechanism of function of *TROLL-8* and demonstrated its involvement in lipid metabolism and its impact on breast cancer progression. Our multi-omics approach, including metabolomic and proteomic analyses, together with the examination of 187 clinical cases and 1064 patient samples in the TCGA breast cancer dataset, provides molecular and functional evidence for the crucial role of *TROLL-8* in FAO. Specifically, we found that *TROLL-8* binds to the FAO rate-limiting enzyme, carnitine palmitoyltransferase 1A (CPT1A), and promotes the interaction between CPT1A and two key components of the FAO pathway, acyl-CoA synthetase long chain family member 1 (ACSL1) and voltage dependent anion channel 1 (VDAC1). We found that *TROLL-8* is required for the production of long-chain fatty acyl-carnitines and affects mitochondrial function and the tumorigenicity of breast cancer cells via a soft agar assay. The physical and functional complex between *TROLL-8*, CPT1A, and its interacting proteins is particularly significant for breast cancers, where we found that high levels of CPT1A and VDAC1 are associated with poor overall survival in all breast cancer patients. Taken together, our data unveil a crucial regulatory mechanism of FAO by *TROLL-8* and demonstrate the importance of the mechanism in tumor progression and survival. Further, these results pave the way for the development of novel therapeutics in breast cancers and other tumors in which FAO has been shown to have a significant role in disease progression.

## RESULTS

### The expression of the TAp63-regulated oncogenic lncRNA-8 (*TROLL-8*) positively correlates with human breast cancer progression

TAp63 has previously been shown to suppress tumor and metastasis formation across multiple tumor types, including breast cancer, through the regulation of miRNA biogenesis,^10^ Hippo pathway signaling,^15^ and oncogenic lncRNA networks.^12^ Importantly, among the lncRNAs we previously found to be regulated by TAp63 in both murine and human breast cancer models,^12^ the expression of TAp63-regulated oncogenic lncRNA-8 (*TROLL-8*) increased along the MCF10A human breast cancer progression model and was highest in the CA1D cell line, which models invasive breast carcinoma.^12^ These RNA-seq results were validated by qRT-PCR, confirming that *TROLL-8* levels are significantly increased between each stage of the MCF10A progression model (Figure 1A). To determine whether this increase in *TROLL-8* levels also occurs in breast cancer patients, we performed RNA *in situ* hybridization (ISH) staining on a tissue microarray (TMA) of breast cancer progression, which included 46 biopsies representing four disease phases: normal breast (NB), lobular hyperplasia (LH), carcinoma *in situ* (CIS), and invasive carcinoma (IC). Notably, quantification of the ISH signal showed a significantly higher expression of *TROLL-8* in tumor biopsies (CIS and IC) compared to normal breast biopsies (Figures 1B and 1C). These findings were further validated using two additional TMAs, totaling 72 NB and 69 IC biopsies, which confirmed that *TROLL-8* levels are elevated in IC compared to NB samples across multiple patient cohorts (Figures 1D-1G). Collectively, these data demonstrate that *TROLL-8* expression increases during breast cancer progression and remains consistently elevated in invasive disease across numerous independent patient groups, supporting its clinical relevance in breast cancer progression.

**Figure 1.**
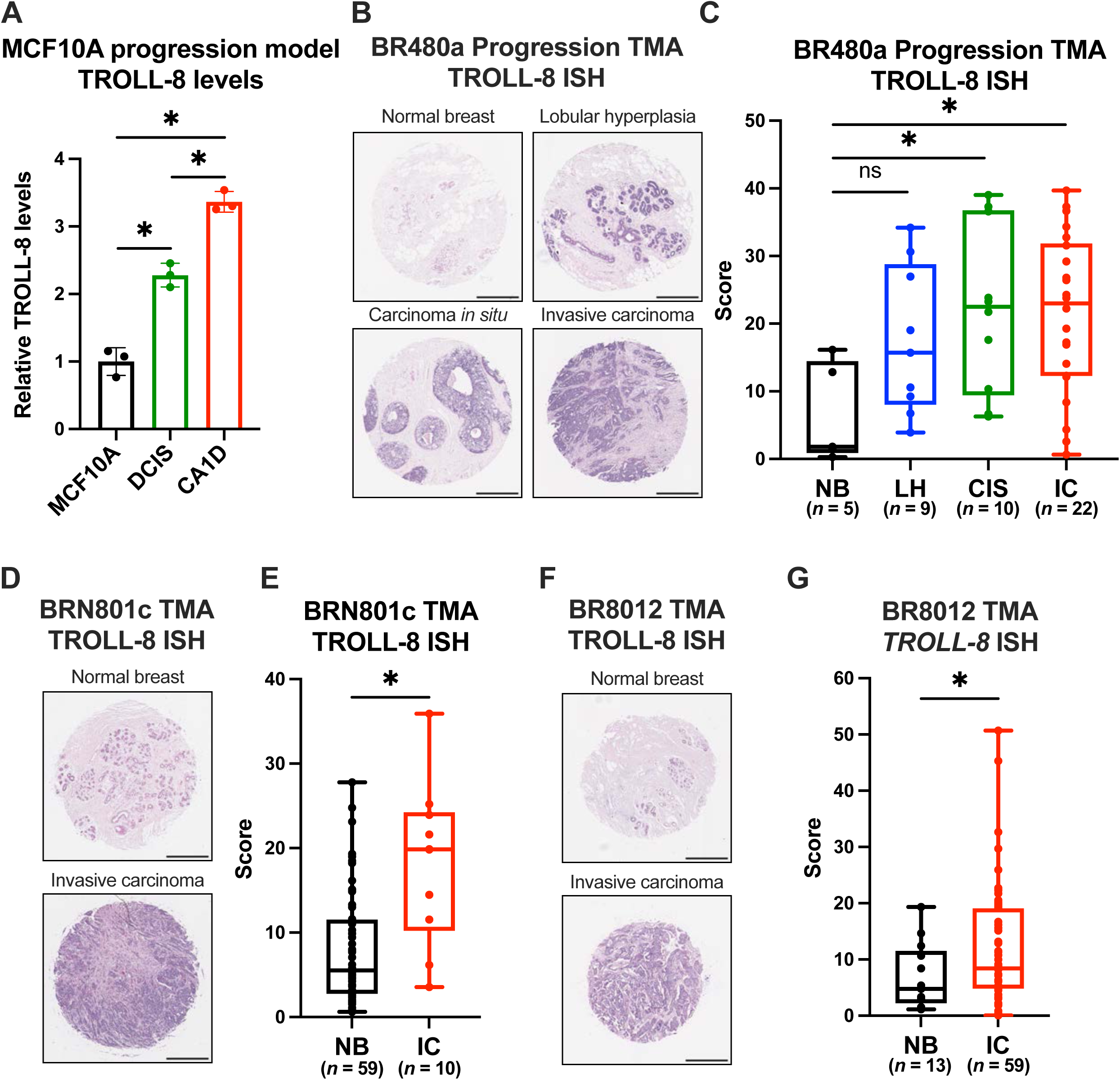
The expression of the TAp63-regulated oncogenic lncRNA-8 (*TROLL-8*) positively correlates with human breast cancer progression. (A) qPCR showing relative *TROLL-8* levels in the human breast cancer progression model. Data represent the mean ± SD of three biological replicates (n = 3). \**p* < 0.05 by Tukey’s multiple comparison test after one-way ANOVA. (B-G) Representative patient tissue microarrays BR480a (B), BRN801c (D), and BR8012 (F) from US Biomax stained for *TROLL-8* expression by ISH with their respective quantifications (C, E, and G). Scale bars represent 500 µm (B, D, and F).All boxplots show the individual data points, the median, interquartile ranges, and the value range (whiskers). \**p* < 0.05 by Dunnett T3 test after Welch ANOVA (C) or Welch’s t-test (E and G).

### *TROLL-8* interacts with proteins enriched in cellular metabolism

LncRNAs typically function by interacting with proteins to regulate various cellular processes, including chromatin remodeling, transcriptional control, RNA stability, and modulation of protein activity signaling.^9,16^ To identify proteins that specifically associate with *TROLL-8*, we hybridized a protein microarray to *in vitro* transcribed (IVT) and fluorescently-labeled RNAs corresponding to the *TROLL-8* sense or antisense strands. Comparison of binding profiles revealed a set of 335 proteins that uniquely interacted with *TROLL-8* but not with its antisense RNA (Figure 2A and Supplementary Table 1). While we found proteins enriched across multiple biological processes (Figure S1A and Supplementary Table 2), more than a quarter of the *TROLL-8* interacting proteins were associated with metabolic pathways, including fatty acid, sugar, amino acid, purine, and NAD metabolism (Figure 2B and Supplementary Tables 3 and 4). Based on the enrichment in metabolic proteins among *TROLL-8* interactors, we next examined whether *TROLL-8* influences cellular metabolism in breast cancer cells. Using the Seahorse mitochondrial stress test assay, we assessed mitochondrial respiration in CA1D cells transfected with either an siRNA pool targeting *TROLL-8* (siTROLL-8) or a non-targeting siRNA (siControl) as a negative control (Figure S1B). Compared to CA1D cells expressing a non-targeting siRNA, *TROLL-8* knockdown decreased mitochondrial respiration, indicating that *TROLL-8* plays a role in maintaining mitochondrial metabolic activity (Figures 2C and 2D). Because mitochondrial respiration can be fueled by multiple metabolic pathways (Figure 2E), we next tested whether *TROLL-8* affects the utilization of specific metabolic substrates by supplementing siControl or siTROLL-8 transfected CA1D cells with glucose, glutamine, or palmitate before measuring mitochondrial respiration. *TROLL-8* knockdown specifically impaired palmitate utilization, while glucose and glutamine metabolism remained unaffected (Figure 2F). These findings suggest that *TROLL-8* interacts with proteins involved in cellular metabolism and influences palmitate-driven mitochondrial respiration.

**Figure 2.**
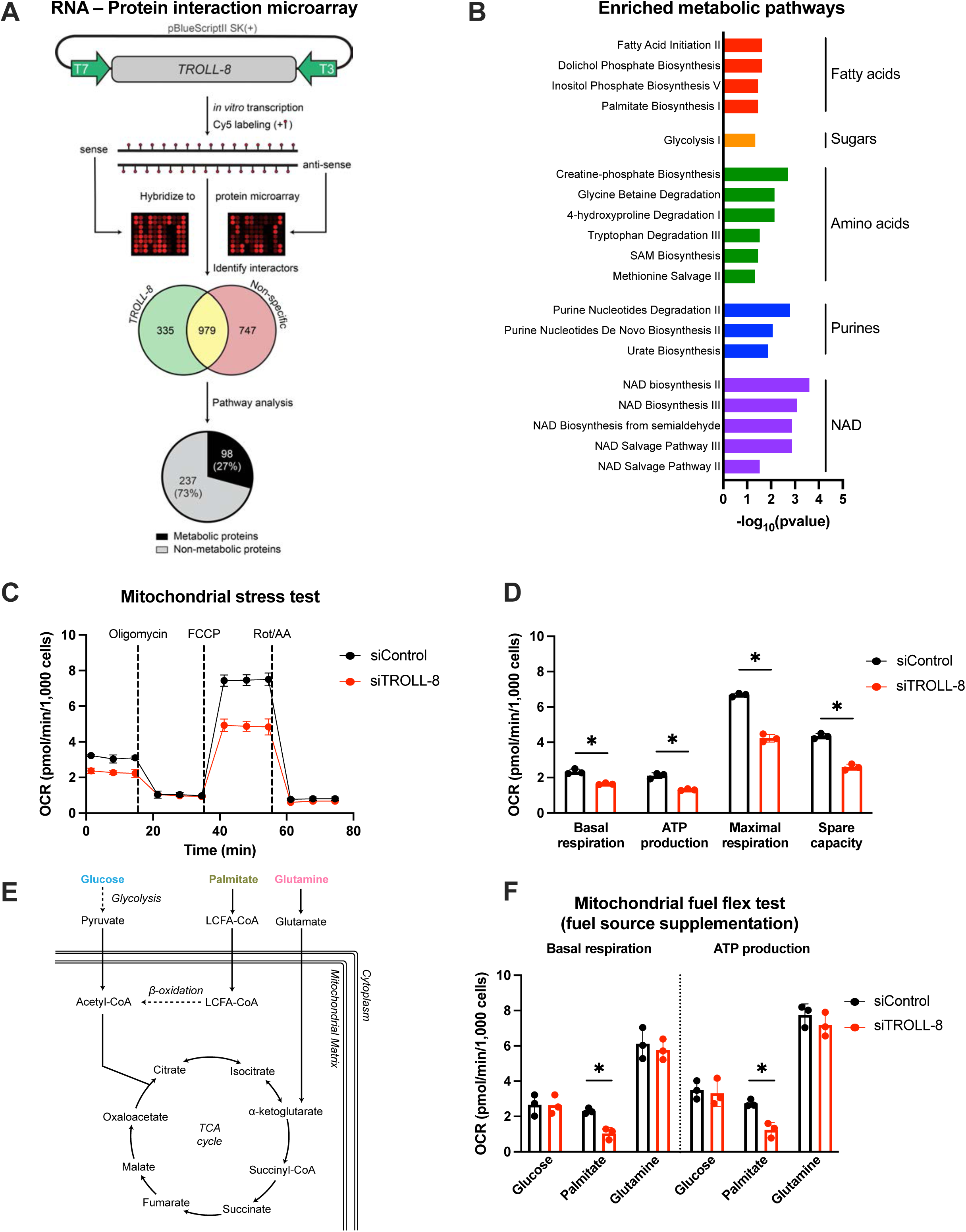
*TROLL-8* interacts with proteins enriched in cellular metabolism. (A) Overview of protein microarray assay, including IVT and labeling of *TROLL-8* and its antisense, test of protein interaction, and pathway analysis of the identified *TROLL-8* interacting proteins. (B) Enriched metabolic pathways among the identified *TROLL-8* interacting proteins. (C) Mitochondrial stress test performed in CA1D cells transfected with siControl (black) or siTROLL-8 (red). (D) Basal respiration, ATP production, maximal respiration, and spare respiratory capacity levels based on the oxygen consumption rate (OCR) values shown in (C). (E) Pathway schematic depicting the uptake of 3 fuel sources: glucose, palmitate, and glutamine. (F) Mitochondrial fuel flex test performed in CA1D cells transfected with siControl (black) or siTROLL-8 (red) and supplemented with the indicated fuel source. Data represent the mean ± SD (n=3). \**p* < 0.05 by unpaired t-test with Holm-Sidak correction for multiple comparisons.

### *TROLL-8* downregulation leads to compromised fatty acid oxidation and the accumulation of long-chain fatty acids

To investigate how *TROLL-8* affects mitochondrial activity, we performed an untargeted metabolomic profiling of CA1D cells transfected with either siTROLL-8 or siControl. Ultra-high-performance liquid chromatography-high resolution mass spectrometry (UHPLC-HRMS) revealed that several long-chain fatty acids (LCFAs) were significantly increased in the *TROLL-8*-depleted cells compared to siControl cells (Figure 3A and Supplementary Table 5), supporting a role for *TROLL-8* in regulating fatty acid metabolism. This accumulation of LCFAs could stem from either increased fatty acid synthesis (FAS) or decreased fatty acid oxidation (FAO). Therefore, to identify which pathway contributed to the observed LCFA enrichment, we performed isotope-tracing experiments by incubating siTROLL-8 and siControl transfected CA1D cells for 48 hours with either U-^13^C-palmitate, U-^13^C-glucose, or U-^13^C-glutamine (Figure 3B). The samples were then analyzed by UHPLC-HRMS to measure the levels of the labeled fuel and the derived isotope-labeled metabolites. Intracellular palmitate levels at the time of collection showed that the amount of U-^13^C-palmitate was higher in the siTROLL-8 samples compared to the siControl counterparts (Figures 3C, 3D, and S2A), indicating that *TROLL-8* knockdown reduces palmitate utilization. This reduced palmitate catabolism was confirmed by the decreased percentage of labeled citrate (Figures 3C, 3E, and S2B), which is produced from the breakdown of palmitate via the β-oxidation pathway. Moreover, we measured the levels of *de novo* synthesized monounsaturated fatty acids (MUFA) and found them to be lower in siTROLL-8 transfected cells compared to siControl transfected cells (Figures 3C, 3F, and S2C). However, tracing U-^13^C-glucose or U-^13^C-glutamine showed that their contributions to the TCA cycle and FAS were unaffected by *TROLL-8* downregulation (Figures 3G-3N and S2D-S2I). Overall, these data indicate that the accumulation of LCFAs caused by *TROLL-8* downregulation results from decreased FAO rather than increased FAS, suggesting that *TROLL-8* may regulate enzymes that control FAO.

**Figure 3.**
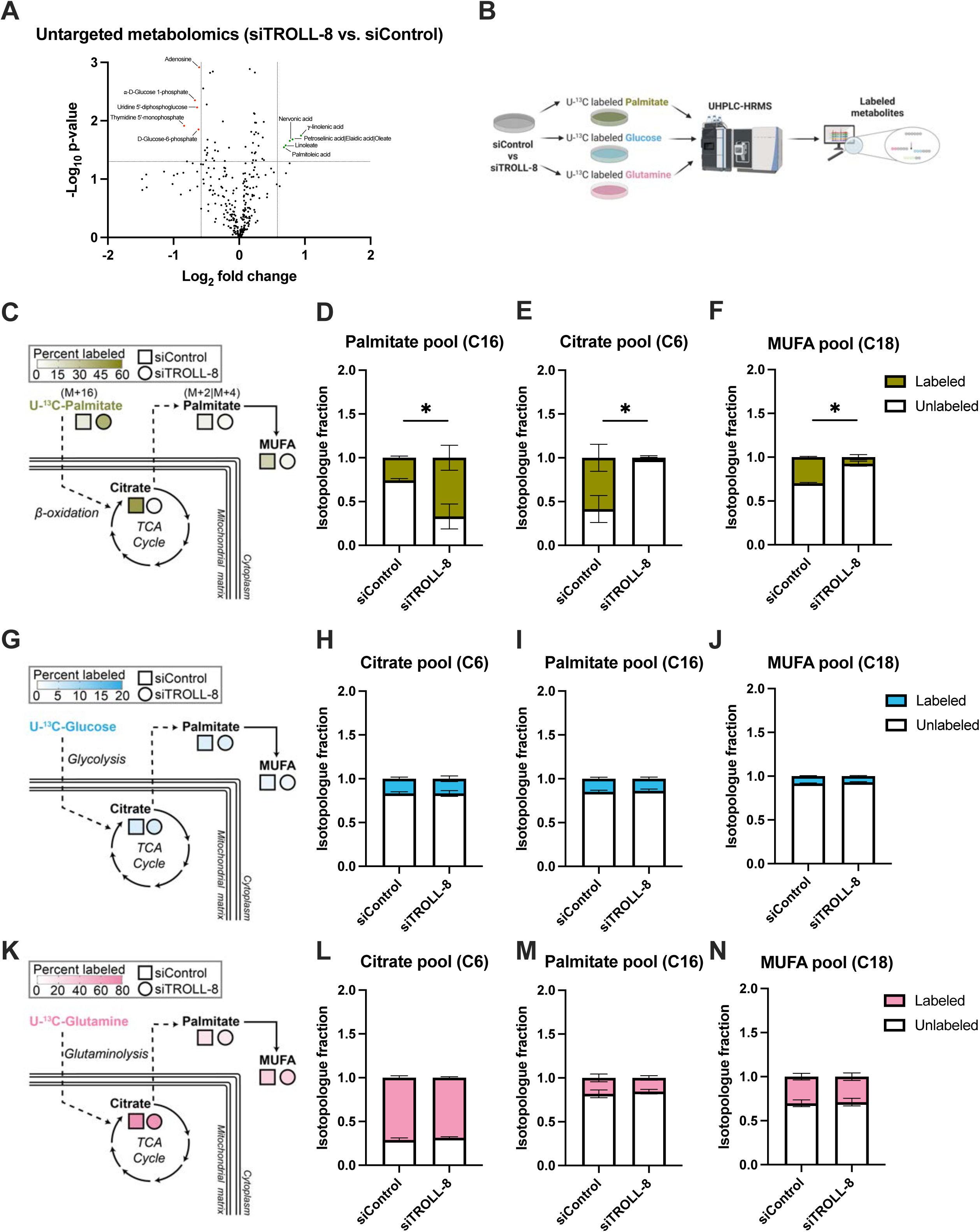
*TROLL-8* downregulation leads to compromised fatty acid oxidation and the accumulation of long-chain fatty acids. (A) Volcano plot representing metabolites that were differentially detected in siTROLL-8 vs. siControl CA1D cells. Metabolites enriched in siTROLL-8 are shown in green, while metabolites depleted in siTROLL-8 are shown in red. Metabolites were considered significantly different if they had a |log_2_(fold change)| > 0.58 and a - log_10_(p-value) > 1.3. (B) Workflow of the isotope tracing experiments via U-^13^C-palmitate, U-^13^C-glucose, or U-^13^C-glutamine in siControl vs siTROLL-8 CA1D cells. (C-F) Schematics depicting the tracing workflow of U-^13^C-palmitate to track the percentage of the indicated labeled metabolites in siControl (square) vs. siTROLL-8 (circle) transfected CA1D cells (C) with respective quantitation of total U-^13^C-palmitate tracing into palmitate (D), citrate (E), and MUFA (F) following culture of CA1D cells transfected with siControl (black) or siTROLL-8 (red) and incubated with U-^13^C-palmitate for 48 hrs. (G-J) Schematics depicting the tracing workflow of U-^13^C-glucose (G) with respective quantitation of total U-^13^C-glucose tracing into citrate (H), palmitate (I), and MUFA (J) following culture of CA1D cells transfected with siControl (black) or siTROLL-8 (red) and incubated with U-^13^C-glucose for 48 hrs. (K-N) Schematics depicting the tracing workflow of U-^13^C-glutamine (K) with respective quantitation of total U-^13^C-glutamine tracing into citrate (L), palmitate (M), and MUFA (N) following culture of CA1D cells transfected with siControl (black) or siTROLL-8 (red) and incubated with U-^13^C-glutamine for 48 hrs. Data represent the mean ± SD (n=3). \**p* < 0.05 for labeled fractions by unpaired t-test.

### *TROLL-8* affects the activity of the FAO rate-limiting enzyme CPT1A

To further elucidate the molecular mechanisms by which *TROLL-8* regulates FAO, we examined our protein microarray screen (Figure 2A and Supplementary Table 1) to identify *TROLL-8*-interacting proteins involved in this pathway. Notably, this led us to focus on carnitine palmitoyltransferase 1A (CPT1A), the rate-limiting enzyme in FAO that is located on the outer mitochondrial membrane and facilitates mitochondrial import of LCFAs for β-oxidation^17^ by joining the fatty acyl-CoAs produced by acyl-CoA synthetase long chain family members (e.g., ACSL1) and free carnitine to form LCF acyl-carnitines (Figure 4A). We first sought to validate the interaction between *TROLL-8* and CPT1A by performing an *in vitro* pull-down assay using the IVT sense and antisense strands of *TROLL-8*. Western blot demonstrated that CPT1A was specifically enriched in the pull-down of the sense transcript (Figure 4B), confirming that indeed CPT1A is a *TROLL-8* interacting protein. Next, we examined whether *TROLL-8* affects CPT1A activity by performing a targeted (acyl-) carnitine metabolomic analysis comparing siControl and siTROLL-8 transfected CA1D cells via UHPLC-HRMS. The targeted carnitine analysis showed enrichment of free carnitine and a simultaneous reduction in LCF acyl-carnitines in *TROLL-8* downregulated cells compared to control cells (Figure 4C), indicating that *TROLL-8* downregulation impaired CPT1A function. To determine if this effect was due to changes in CPT1A protein levels or localization in the absence of *TROLL-8*, we performed cellular fractionation assays. As shown in Figure 4D, neither CPT1A protein levels nor its mitochondrial localization were affected by the downregulation of *TROLL-8*. Additionally, CPT1A mRNA levels were unaltered too (Figures S3A and S3B). Together, these results indicate that *TROLL-8* regulates FAO by directly binding to CPT1A and regulating its enzymatic activity.

**Figure 4.**
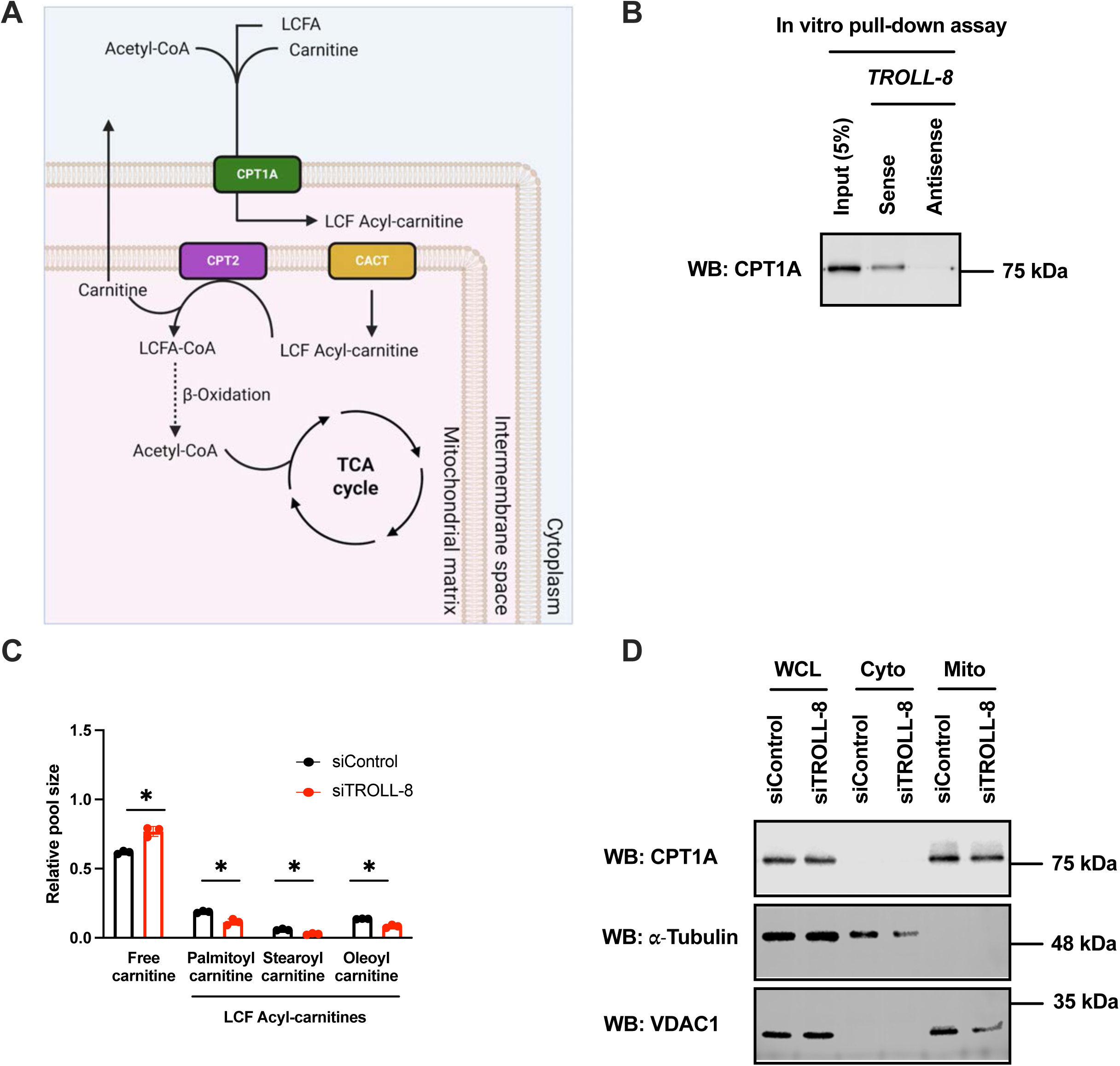
*TROLL-8* affects the activity of the FAO rate-limiting enzyme CPT1A. (A) Diagram depicting the FAO pathway. (B) Representative western blot for CPT1A pulled down by the biotinylated *TROLL-8* sense or antisense strand. (C) Targeted metabolomics quantifying the indicated metabolites in siControl (black) vs siTROLL-8 (red) transfected cells. Data represent the mean ± SD (n=3). \**p* < 0.05 by unpaired t-test with Holm-Sidak correction for multiple comparisons. (D) Representative western blot for CPT1A in the whole cell lysate (WCL), cytoplasmic (Cyto), and mitochondrial (Mito) fractions of CA1D cells transfected with siControl or siTROLL-8.

### The CPT1A-ACSL1-VDAC1 complex mediates the pro-oncogenic activities of TROLL-8

To explore how *TROLL-8* regulates the enzymatic activity of CPT1A, we immunoprecipitated CPT1A in CA1D cells transfected with siControl or siTROLL-8 and conducted ultra high-performance liquid chromatography tandem mass spectrometry (LC-MS/MS) analysis to identify proteins whose interaction with CPT1A is affected by *TROLL-8* (Figure 5A and Supplementary Table 4). Among the identified proteins whose interaction with CPT1A decreased with *TROLL-8* downregulation, ACSL1 and VDAC1 were particularly notable due to their essential roles in FAO and localization to the outer mitochondrial membrane.^18,19^ To confirm the LC-MS/MS proteomics results, we performed an immunoprecipitation for CPT1A in CA1D cells transfected with either siControl or siTROLL-8 and observed that the interaction of CPT1A with ACSL1 and VDAC1 indeed diminishes in the absence of *TROLL-8* (Figure 5B). To determine if the loss of the CPT1A-interacting proteins mimics the mitochondrial defects seen with *TROLL-8* depletion (see Figures 2C and 2D), we evaluated mitochondrial function following individual knockdown of *TROLL*-8, ACSL1, CPT1A, or VDAC1 (Figures S4A-S4D). Notably, depletion of ACSL1, CPT1A, or VDAC1 each phenocopied the mitochondrial respiration defects observed with *TROLL-8* knockdown (Figures 5C and S4E-S4H). Indeed, their downregulation caused a significant reduction in mitochondrial oxygen consumption rate compared to the control cells, supporting a functional connection between *TROLL-8* and the CPT1A-associated FAO proteins.

**Figure 5.**
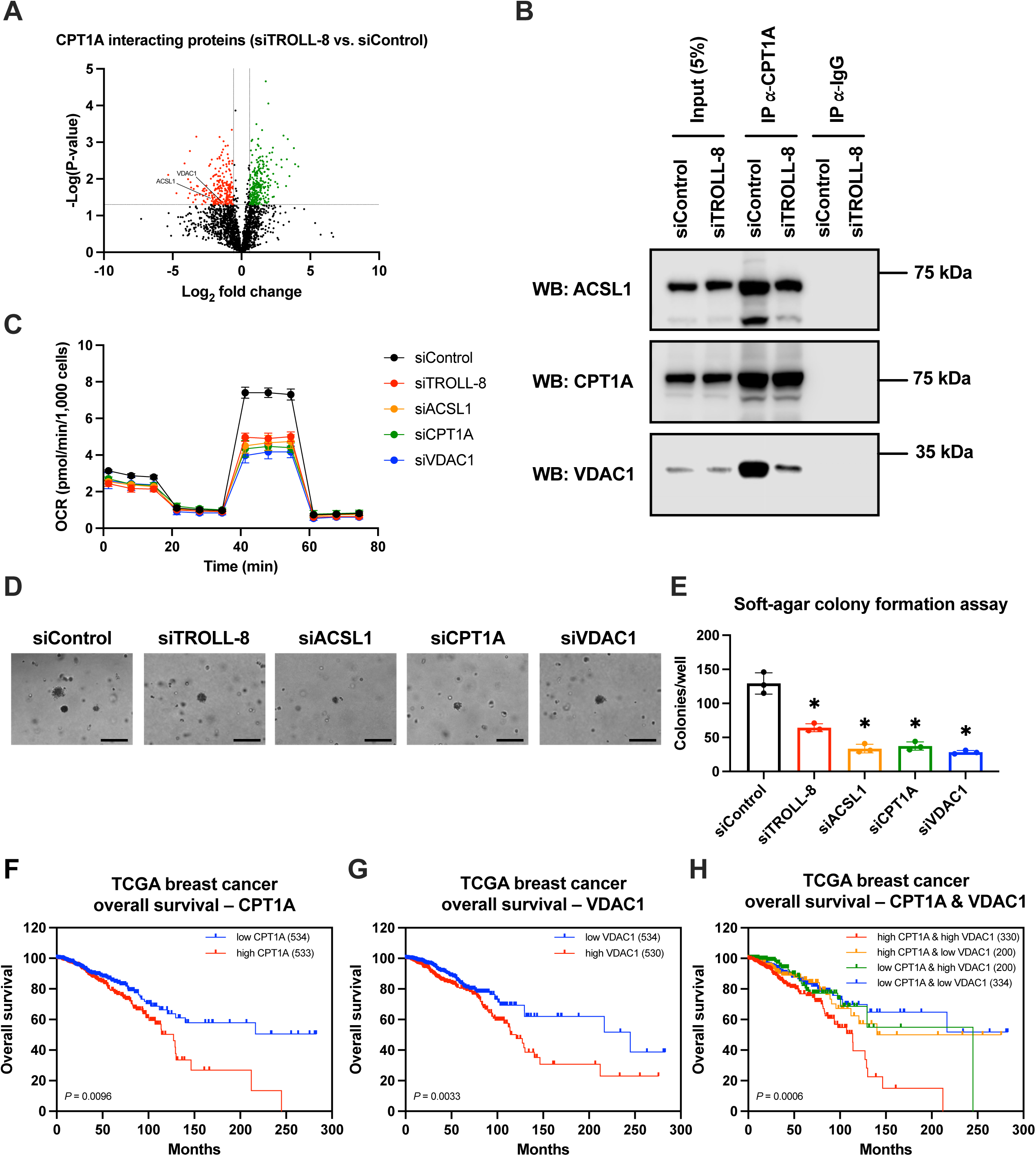
The CPT1A-ACSL1-VDAC1 complex mediates the pro-oncogenic activities of *TROLL-8*. (A) Volcano plot representing proteins that were differentially bound to CPT1A in siTROLL-8 vs. siControl transfected CA1D cells. Proteins whose interaction with CPT1A is significantly increased or decreased in siTROLL-8 are shown in green and red, respectively. Changes in interactions were significant if they had a |log_2_(fold change)| > 0.58 and a - log_10_(p-value) > 1.3. (B) Representative western blot analysis of the immunoprecipitation of endogenous CPT1A in CA1D cells transfected with the indicated siRNAs. (C) Mitochondrial stress test performed in CA1D cells transfected with the indicated siRNAs. (D) Representative image of soft agar colonies of CA1D cells transfected with the indicated siRNAs. Scale bars represent 200 µm. (E) Quantification of soft agar assay colonies in CA1D cells transfected with the indicated siRNAs. Data represent, mean ± SD (n=3). \**p* < 0.05 by Dunnett’s multiple comparisons test against siControl after one-way ANOVA. (F-H) Kaplan-Meier curves of overall breast cancer survival data from the TCGA Invasive Breast Carcinoma GDC 2025 dataset, stratified by the expression of CPT1A (F), VDAC1 (G), or both (H). P-values were calculated using the log-rank test via cBioPortal.

Because the role of metabolic reprogramming has been linked to promoting breast cancer progression,^20^ we next assessed whether mitochondrial defects linked to the loss of *TROLL-8* and the CPT1A-interacting proteins influence tumorigenic potential. To measure anchorage-independent growth, a hallmark of cellular transformation,^21^ we performed soft agar colony formation assays after knocking down *TROLL-*8, ACSL1, CPT1A, or VDAC1 individually. Downregulation of any of these factors significantly impaired anchorage-independent growth compared to the control cells (Figures 5D and 5E), indicating that disrupting mitochondrial FAO limits the tumorigenic potential of CA1D breast cancer cells. To evaluate the clinical relevance in human breast cancer patients of these FAO-related proteins, we analyzed the breast cancer dataset from The Cancer Genome Atlas (TCGA).^22^ While ACSL1 expression did not significantly stratify patient outcomes, possibly due to functional redundancy among ACSL family members,^23^ higher expression of CPT1A or VDAC1 correlated with poorer overall survival (Figures 5F and 5G). These findings suggest a clinically important role for CPT1A and VDAC1 in breast cancer progression and align with our functional data, showing that these proteins partner with *TROLL-8* to support mitochondrial FAO and tumor growth. We then examined patient outcomes based on combined CPT1A and VDAC1 expression. Survival analysis revealed that patients with high expression of both CPT1A and VDAC1 had the lowest overall survival (Figure 5H), consistent with the idea that efficient long-chain fatty acid import and active mitochondrial FAO may promote more aggressive disease. Conversely, patients with low expression of one or both genes, indicating decreased LCFA import into the mitochondria, had improved overall survival. Overall, these findings demonstrate that *TROLL-8* regulates CPT1A activity by facilitating its interaction with ACSL1 and VDAC1, thereby promoting FAO and mitochondrial respiration (Figure 6). We also show that the regulation of CPT1A by *TROLL-8* contributes to the tumorigenic potential of breast cancer cells and worse overall survival in breast cancer patients.

**Figure 6.**
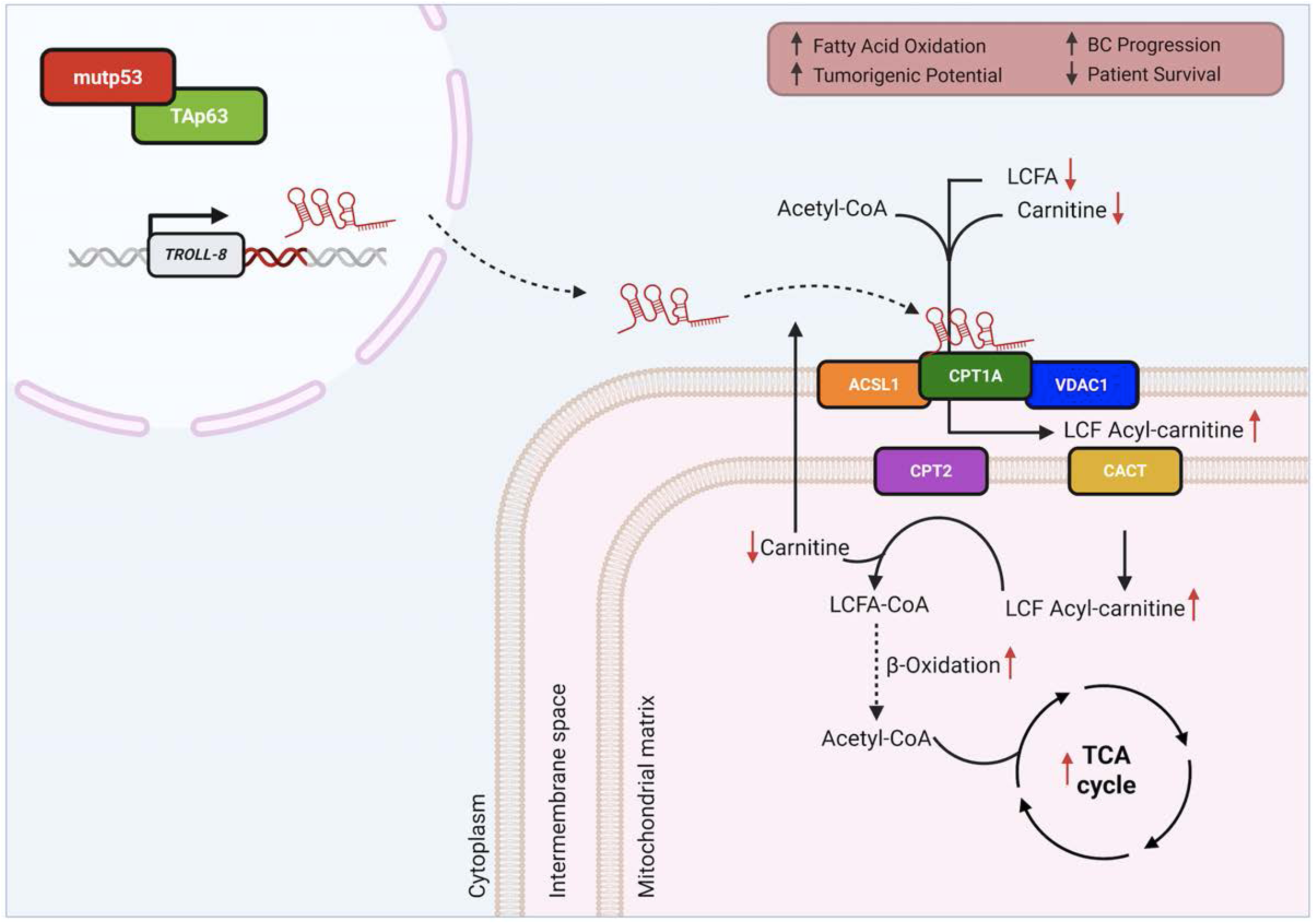
Molecular mechanism of *TROLL-8* function. Model depicting the role of *TROLL-8*. In tumors expressing mutant p53, the functions of the tumor and metastasis suppressor, TAp63, are inhibited. This leads to the expression of *TROLL-8*. This lncRNA binds to CPT1A, allowing CPT1A to interact with ACSL1 and VDAC1, thus leading to increased synthesis and oxidation of LCF acyl-carnitines (indicated by the red arrows). This leads to an increase in mitochondrial respiration via the TCA cycle, increased tumorigenic potential of breast cancer cells, and poor overall survival in breast cancer patients.

## DISCUSSION

Metabolic reprogramming is a hallmark of cancer, enabling cancer cells to survive, proliferate, and resist therapies by rewiring nutrient uptake and utilization, ultimately driving cancer progression.^24^ Increasing evidence has recently shown that lipid metabolism is altered in multiple tumor types due to changes in mitochondrial function and FAO, fueling aggressive cancer states.^3,4^ We previously demonstrated that the tumor and metastasis suppressor TAp63, a member of the p53 family of transcription factors, regulates metabolism^11^ and a network of conserved lncRNAs.^12^ Our current findings connect these areas by demonstrating that a TAp63-regulated lncRNA, *TROLL-8*, is a key regulator of FAO that promotes breast cancer progression. Indeed, we found that *TROLL-8* binds to the FAO rate-limiting enzyme, carnitine palmitoyltransferase 1A (CPT1A), and promotes the interaction of CPT1A with acyl-CoA synthetase long chain family member 1 (ACSL1) and voltage dependent anion channel 1 (VDAC1), thereby enhancing fatty acid catabolism, increasing tumorigenic potential, and advancing breast cancer progression, ultimately impacting the survival of breast cancer patients (Figure 6).

Beyond its role in energy production, FAO has been linked to multiple cancer hallmarks. Indeed, FAO regulates redox homeostasis via NADPH production,^25^ maintains mitochondrial health,^26^ and enables tumor cells to adapt to stressful environments.^27^ Additionally, FAO has been shown to be involved in stem cell states,^28^ immune evasion,^29^ and metastatic spread in various tumor types.^30^ Specifically, in the case of breast cancers, FAO has been associated with increased stemness and proliferation due to the upregulation of CPT1A driven by low CD24 expression.^31^ Given that CD24-low tumors comprise about one-third of breast cancer cases, especially aggressive IC,^32^ which we found tend to have high *TROLL-8* expression, this mechanism may contribute to FAO-dependent programs across a broad subset of patients.

Our analysis of the breast cancer TCGA dataset revealed that patients with high CPT1A and VDAC1 levels have the poorest overall survival, highlighting the clinical significance of this axis. Since elevated CPT1A^30^ and increased FAO^33^ have been associated with resistance to chemotherapy and radiation therapy,^34^ several approaches targeting FAO via CPT1A inhibition have been explored. These treatment strategies include irreversible covalent inhibitors that bind to the active site of CPT1A, such as etomoxir and 2-tetradecylglycidic acid (TDGA),^17^ as well as irreversible competitive inhibitors of CPT1A, such as oxfenicine.^35^ However, their clinical application has been limited by toxicity, prompting the exploration of alternative strategies that instead target CPT1A interactions with other proteins. Intriguingly, one of these is the small molecule DHP-B, which induces apoptosis by disrupting the interaction between CPT1A and VDAC1,^36^ and could be especially effective in tumors with high TROLL-8 levels that may rely on a stronger interaction between CPT1A and VDAC1. Likewise, direct knockdown of TROLL-8, which is highly upregulated in invasive breast cancer, or disruption of its interaction with CPT1A, could offer a more selective approach for disrupting FAO that is less likely to induce the toxicity seen with many small-molecule inhibitors.

In conclusion, our findings provide new mechanistic insights into the connections between FAO and breast cancer progression through our complementary methods, including the analysis of breast cancer tissue microarrays, the integration of metabolomics and proteomics profiles, and the molecular and functional study of *TROLL-8* and its interacting proteins. These results demonstrate that this lncRNA is essential for the effective binding of CPT1A to ACSL1 and VDAC1, which then influences mitochondrial respiration and the tumorigenic potential of breast cancer cells (Figure 6), as well as patient outcomes. Finally, these findings suggest new possibilities for therapies targeting *TROLL-8* and other components of this pathway, which could be especially effective in breast cancer and potentially other tumor types where FAO plays a significant pro-oncogenic role.

## Supporting information

Supplementary Table 1_TROLL-8 sense and antisense interacting proteins

Supplementary Table 2_Non-metabolic pathways of TROLL-8 interacting proteins

Supplementary Table 3_Metabolism-associated proteins interacting with TROLL-8

Supplementary Table 4_Metabolic pathways of TROLL-8 interacting proteins

Supplementary Table 5_Untargeted metabolomics in siControl vs siTROLL-8

Supplementary Table 6_CPT1A interacting proteins identified by mass spec

## SUPPLEMENTARY FIGURE LEGENDS

**Figure S1:**
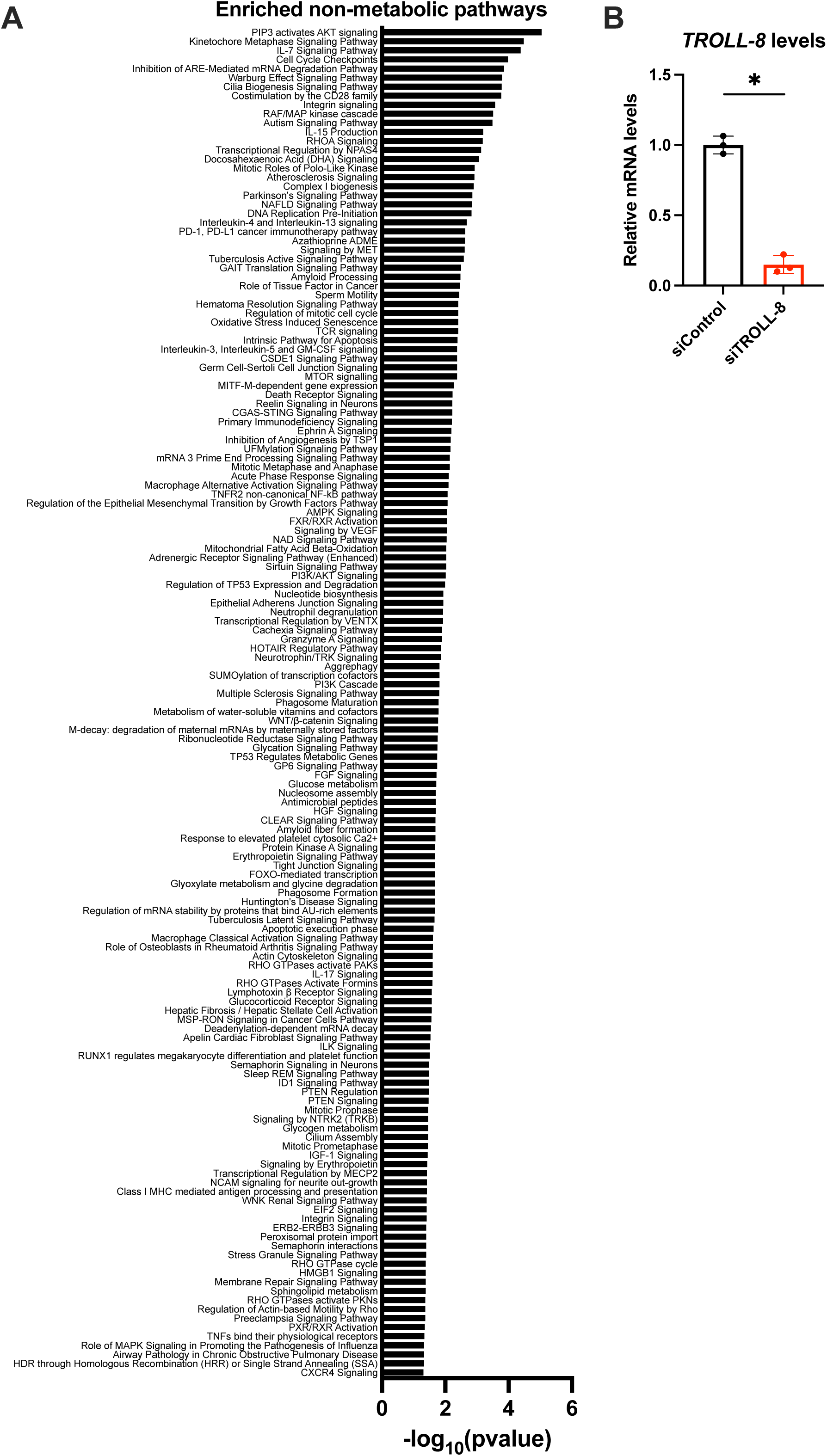
*TROLL-8* interacts with proteins enriched in cellular metabolism. (A) Enriched non-metabolic pathways among the identified *TROLL-8* interacting proteins. (B) qPCR of *TROLL-8* mRNA levels in CA1D cells transfected with siRNAs. Data represent mean ± SD (n=3). \**p* < 0.05 by unpaired t-test.

**Figure S2:**
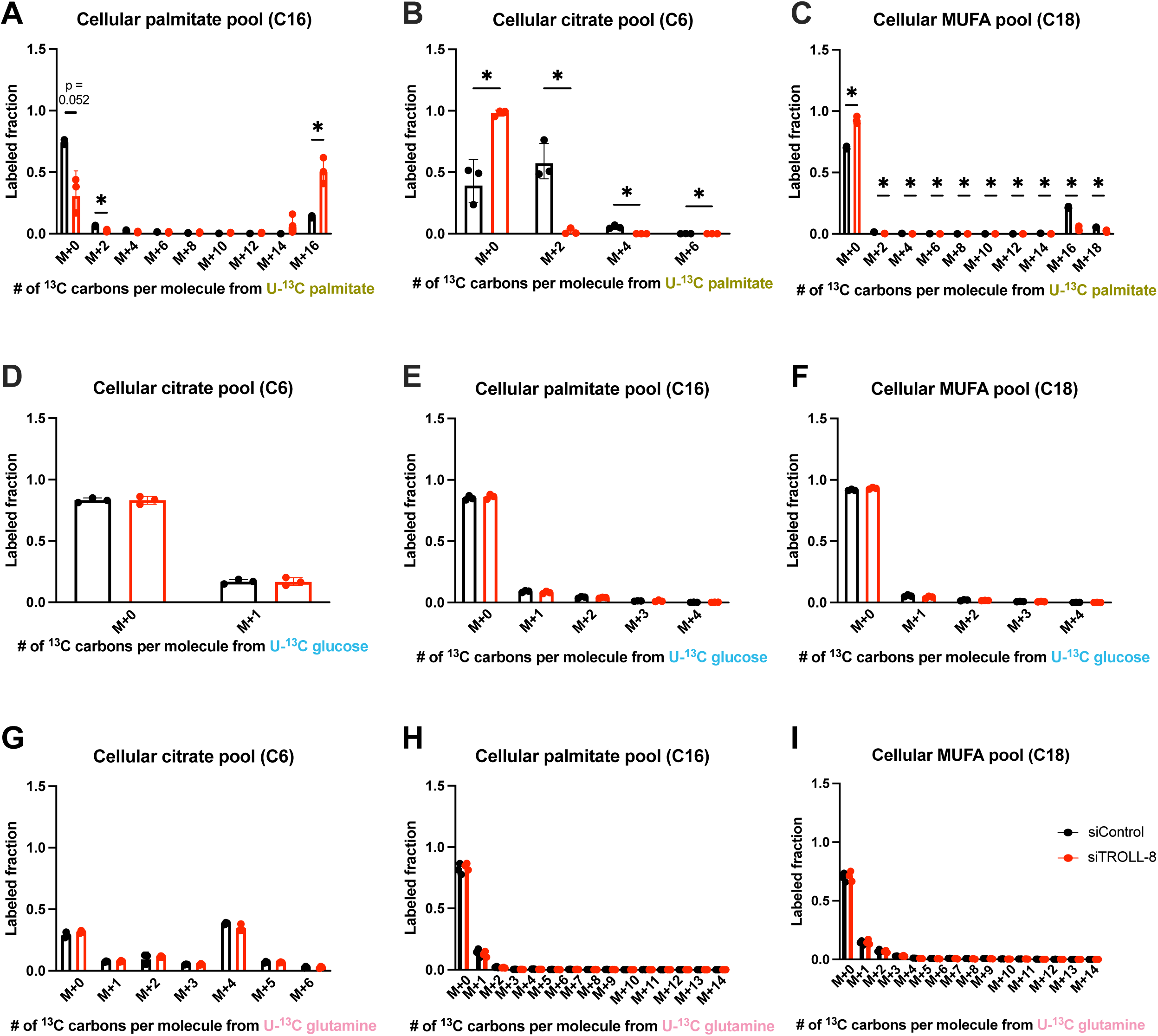
*TROLL-8* downregulation leads to compromised fatty acid oxidation and the accumulation of long-chain fatty acids. (A-C) Quantitation of U-^13^C-palmitate tracing into palmitate (A), citrate (B), and MUFA (C) following culture of CA1D cells transfected with siControl (black) or siTROLL-8 (red) and incubated with U-^13^C-palmitate for 48 hrs. (D-F) Quantitation of U-^13^C-glucose tracing into citrate (D), palmitate (E), and MUFA (F) following culture of CA1D cells transfected with siControl (black) or siTROLL-8 (red) and incubated with U-^13^C-glucose for 48 hrs. (G-I) Quantitation of U-^13^C-glutamine tracing into citrate (G), palmitate (H), and MUFA (I) following culture of CA1D cells transfected with siControl (black) or siTROLL-8 (red) and incubated with U-^13^C-glutamine for 48 hrs. Data represent the mean ± SD (n=3). \**p* < 0.05 by unpaired t-test with

**Figure S3:**
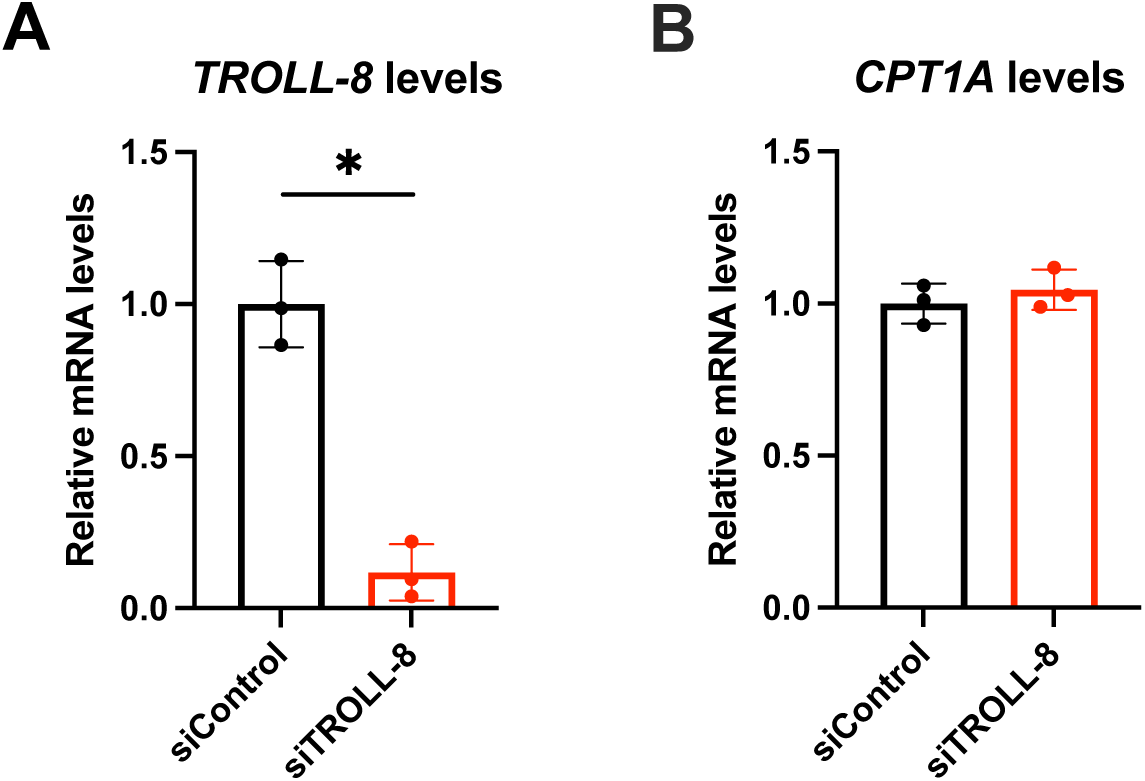
*TROLL-8* affects the activity of the FAO rate-limiting enzyme CPT1A. (A-B) qPCR of *TROLL-8* (A) and *CPT1A* (B) mRNA levels in CA1D transfected with the indicated siRNAs. Mean ± SD (n=3). Data were analyzed with Welch’s t test, \**p* < 0.05.

**Figure S4:**
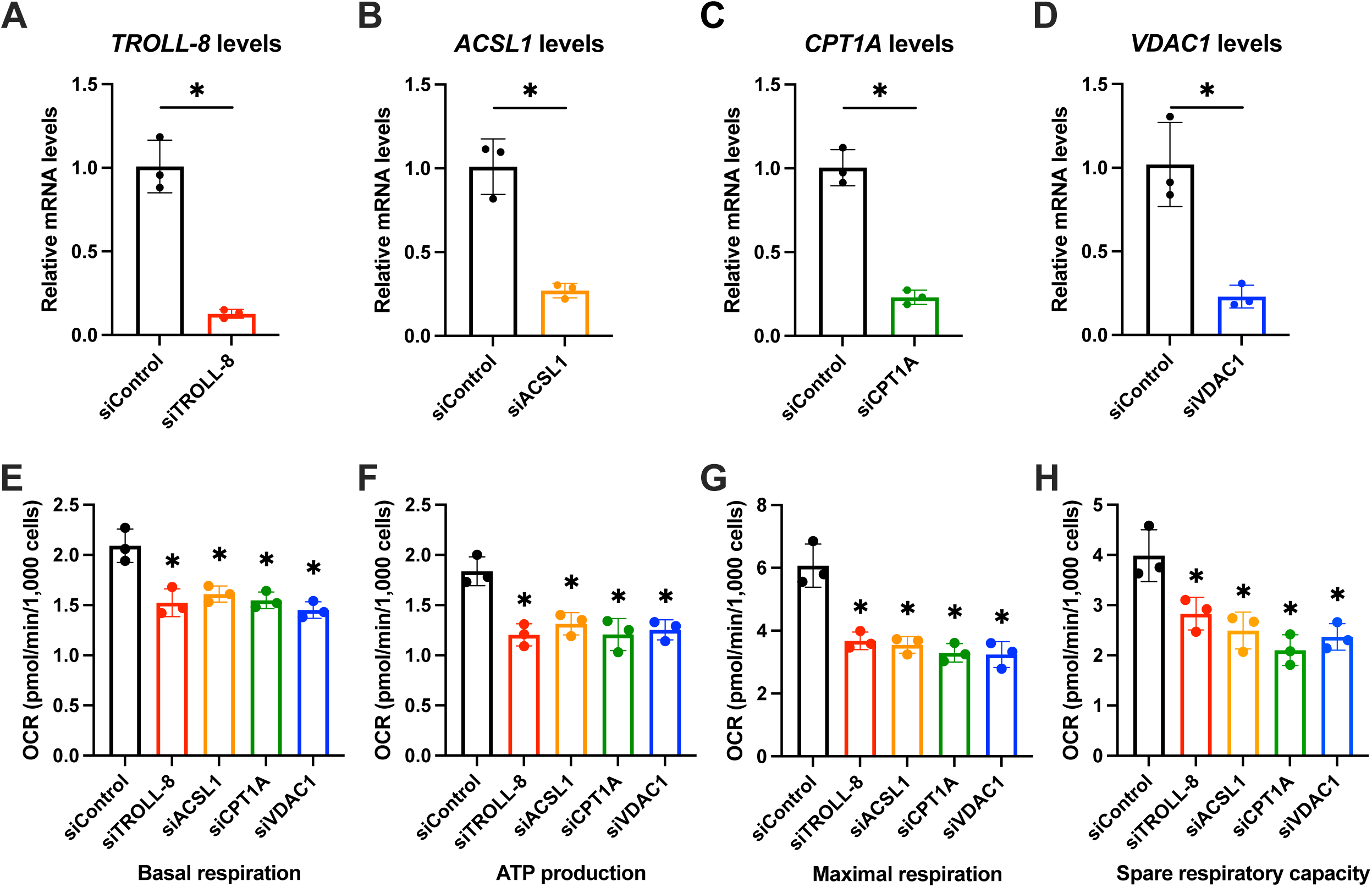
The CPT1A-ACSL1-VDAC1 complex mediates the pro-oncogenic activities of *TROLL-8*. (A-D) qPCR of the indicated mRNAs in CA1D cells transfected with either siControl and siTROLL-8 (A), siACSL1 (B), siCPT1A (C) or siVDAC1 (D). (E-H) Basal respiration (E), ATP production (F), maximal respiration (G), and spare respiratory capacity (H) levels based on the oxygen consumption rate (OCR) values shown in Figure 5C. Mean ± SD (n=3). Data were analyzed with Welch’s t test (A-D) and one-way ANOVA (E-H), \**p* < 0.05.

## RESOURCE AVAILABILITY

### Lead contact

Requests for further information and resources should be directed to and will be fulfilled by the lead contact, Elsa R. Flores.

### Materials availability

All unique/stable reagents generated in this study are available from the lead contact without restriction.

### Data and code availability

All data and materials reported in this paper will be shared upon request by the lead contact. This paper does not report original code.

Proteomics data have been deposited to the ProteomeXchange Consortium via the PRIDE^37^ partner repository with the dataset identifier PXD078984 and 10.6019/PXD078984.

Metabolomics data have been uploaded to MetaboLights^38^ repository with the study identifiers: MTBLS14632 (Untargeted Metabolomics), MTBLS14721 (Isotope Tracer Experiments), and MTBLS14731 (Acyl-Carnitines).

## ACKNOWLEDGEMENTS

We thank Joseph Johnson and his team at the Analytic Microscopy Core Facility (Moffitt Cancer Center) for the quantification of the ISH signals.

This work was supported by the National Cancer Institute (R35-CA197452 and P01-CA250984), the Florida Cancer Innovation Fund (MOAZZ), and a V Foundation All Star Translational Award (AST2026-009) to E.R.F.

J.H.L. was supported by the Integrated Program in Cancer and Data Science postdoctoral training fellowship (T32-CA233399).

This work has been supported in part by the Proteomics and Metabolomics Core Facility at the H. Lee Moffitt Cancer Center & Research Institute, an NCI-designated Comprehensive Cancer Center (P30-CA076292).

We thank BioRender (www.biorender.com) for assistance with the creation of figure illustrations.

## AUTHOR CONTRIBUTIONS

A.A.D., M.N., and E.R.F. conceived the study, designed experiments, and analyzed data. A.A.D., M.N., and X.L. designed and performed experiments. A.A.D., M.N., J.H.L., X.L., and E.R.F. analyzed the data. A.A.D., M.N., and J.H.L. performed data visualization. M.L. and J.M.K. ran and analyzed the metabolomics experiments. L.N.F.D. and J.M.K. ran and analyzed the proteomics experiments. A.A.D., M.N., J.H.L., and E.R.F. wrote the paper. All authors discussed and approved the paper.

## METHODS

### *In situ* hybridization of human tissue microarrays

TMAs of breast cancer progression (US Biomax, BR480a) and of breast normal adjacent tissue and cancer tissue (US Biomax, BRN801c and BR8012) were used for the ISH assay. The ISH was performed using the miRCURY LNA miRNA kit for FFPE (Qiagen, 339450), as previously reported^12^. The following probe was utilized to detect *TROLL-8* at 55°C for 1 h: 5’ – TACAGAGGCAAGCGGTGAACT – 3’. The signal intensity (continuous variable, 0–1) and the proportion of positive tissue (continuous variable, 0–100%) were quantified using the Oncotopix® software (Visiopharm) and subsequently multiplied to obtain a value comprised between 0 and 100, which represents the ISH score.

### Cell lines and culture conditions

MCF10A, DCIS, and CA1D cells were purchased from the Karmanos Cancer Institute (Detroit, MI) and grown in DMEM/F12 (1:1) supplemented with 5% horse serum (Gibco, 16050122), 1% Penicillin-Streptomycin (ThermoFisher Scientific, 15-140-122), 20 ng/mL human epidermal growth factor (Gibco/PeproTech, 10781-692), 10 µg/mL insulin (Sigma Aldrich, I9278), 500 ng/mL hydrocortisone (Sigma-Aldrich, H0888-1G). All cultured cells were maintained at 37°C with 5% CO_2_ and tested periodically for mycoplasma using the MycoAlert Kit (Lonza, LT07-710).

### Gene expression analysis by quantitative real time PCR

Total RNA was prepared using the miRNeasy Mini kit (QIAGEN, 217004). For gene expression analysis, complementary DNA was synthesized using the SuperScript II First-Strand Synthesis Kit (Invitrogen, 18064-014) according to the manufacturer’s protocol followed by qRT-PCR using the TaqMan® Universal PCR Master Mix (Applied Biosystems, 4364343). qRT-PCR was performed in triplicate using the QuantStudio 6 flex PCR machine (Applied Biosystems). The following Taqman assays were used to detect the indicated genes: Hs00960561_m1 (ACSL1), Hs00912671_m1 (CPT1A), Hs00172187_m1 (POLR2A), Hs00922963_s1 (*TROLL-8*), and Hs04978484_m1 (VDAC1).

### Protein microarray analysis of Cy5-labeled *TROLL-8* followed by Ingenuity Pathway Analysis

*In vitro* transcription of the sense and antisense strands of *TROLL-8* RNA was performed from the pBluescript II SK (+) – *TROLL-8* vector using T7 and T3 polymerase (MEGAscript^TM^ T7/T3 Transcript Kits, Thermo Fisher Scientific, AM1334/ AM1338) in accordance with the manufacturer’s instructions. Following transcription, the strands were labelled with Cy5 using the Label IT µArray Cy5 labelling kit (Mirus, MIR 3700) with a labelling efficiency of 3 pmol Cy5 dye per µg of RNA. Protoarray Human Protein Microarrays v5.2 (Thermo Fisher Scientific, PAH0525101) were used for the hybridization with 10 pmol of either Cy5 labelled sense or antisense as a negative control. Three independent replicates were carried out with the hybridization step performed at 4 °C for 1 h in the dark.^12^ The slides were then scanned with the GenePix Microarray Scanner (Molecular Devices, 4000B) at 635 nm to select proteins with mean signal intensities greater than 2 (F635/B635 > 2) in all three replicates. The resulting proteins were then categorized based on their interaction with either the sense strand of *TROLL-8*, its antisense strand, or both (see Supplementary Table 1). The proteins interacting exclusively with the sense strand of *TROLL-8* were then analyzed using PantherDB GO-Slim Biological Process^39,40^ to identify enriched pathways and biological processes. The metabolism-associated proteins (“metabolic process – GO:0008152”) interacting with *TROLL-*8 are listed in Supplementary Table 3. Pathway analysis of the full set of 335 *TROLL-8* interacting proteins was performed using Ingenuity Pathway Analysis (Qiagen) and divided into non-metabolic and metabolic pathways (Supplementary Tables 2 and 4).

### siRNA sequences and transfection

For siRNA transfection, double-stranded RNA oligos (40nM) were transfected using Lipofectamine RNAiMAX (Invitrogen) followed 24 h afterwards by Lipofectamine 2000 (Invitrogen) according to the manufacturer’s instructions. The universal negative siRNA control #1 (siControl) was purchased from Sigma-Aldrich (SIC001-10NMOL). Additional siRNAs utilized were: siACSL1 (SASI_Hs01_00202187, Sigma), siCPT1A (5’ – CCCUAAAUGUAAGGGAGAU – 3’), siTROLL-8 pool^12^, and siVDAC1 (SASI_Hs01_00012464, SASI_Hs01_00012466, SASI_Hs01_00012467, Sigma).

### Seahorse assay

For each biological replicate, CA1D cells were transfected with the indicated siRNAs and then seeded at 3.0 x 10^4^ cells/well (6 technical replicates/each) in a 96-well Agilent Seahorse XF Cell Culture Microplate (Agilent, 103794-100). The oxygen consumption rate (OCR) of the cells was measured by the Seahorse XF96 extracellular flux analyzer (Agilent Technologies) using a Seahorse XF Cell Mito Stress Test kit (Agilent Technologies, 103015-100) in accordance with the manufacturer’s instructions. For glucose and glutamine supplementation studies, the assay medium was prepared by supplementing the prewarmed Seahorse XF base medium with 5 mM HEPES and either 10 mM glucose or 2mM glutamine. For the fatty acid supplementation assay, the assay medium was prepared by supplementing 1x KHB buffer (111 mM NaCl, 4.7 mM KCl, 1.25 mM CaCl_2_, 2.0 mM MgSO_4_, and 1.2 mM NaH_2_PO_4_) with 2.5 mM glucose, 0.5 mM carnitine, and 5 mM HEPES.

### Untargeted metabolomics

#### Sample preparation

CA1D cells were transfected with either siControl or siTROLL-8. 48 h post-transfection, cells were harvested and washed 2-3 times with ice-cold PBS. The sample collection and the subsequent processing were carried out on ice. An aliquot of the internal standard mixture was added to each sample, and metabolites were extracted by adding 1mL of precooled 80% methanol (stored in -80 °C for at least one hour prior to extraction) to precipitate proteins. Following the addition of the extraction solvent, the samples were vortexed and centrifuged. Samples were then incubated at -80 °C for 30 min to increase metabolite extraction. After incubation, the samples were immediately centrifuged, and the supernatant was transferred to a new microcentrifuge tube. The protein pellet was resolubilized using aqueous 20 mM HEPES with 8 M urea for Bradford assays to measure the protein concentration. Dried metabolites were re-dissolved in 20 μL aqueous 80% methanol.

#### UHPLC-HRMS metabolomics

Ultra high-performance liquid chromatography-high resolution mass spectrometry (UHPLC-HRMS) was performed using a Vanquish UHPLC interfaced with a Q Exactive HF quadrupole-orbital ion trap mass spectrometer (Thermo Fisher Scientific). Chromatographic separation was performed using a SeQuant ZIC-pHILIC guard column (2.1 mm ID x 20 mm length, 5 µm particle size) and a SeQuant ZIC-pHILIC LC column (2.1 mm ID x 150 mm length, 5 µm particle size, MilliporeSigma). The column was kept in a 30 °C column chamber and 2 µL of sample loaded via autosampler. For the gradient, mobile phase A (10 mM ammonium carbonate and 0.05% ammonium hydroxide) and mobile phase B (100% acetonitrile) were used as follows (total run time 20 min): 0-13 min, 80% to 20% of B, 13-15 min, 20% of B, 15-20 min 80% of B. Full MS was performed in positive and negative mode separately detecting ions from m/z 65-900.

#### Reagents and chemicals

Ammonium hydroxide and ammonium carbonate were obtained from MilliporeSigma. LC-MS grade solvents, including water, methanol and acetonitrile, were purchased from Honeywell Burdick and Jackson (sourced through VWR). The internal standards were obtained from Cambridge Isotope Labs.

#### Metabolomics data analysis

MZmine software, version 3.39, was used to identify and quantify metabolites by matching by m/z and RT to an in-house library. Data normalization was performed based on protein concentration. The identified metabolites, relative fold changes, and p-values are listed in Supplementary Table 3.

### Untargeted metabolomics of tracing experiments

For the palmitate tracing experiments, the transfected cells were incubated for 72 h with DMEM/F-12 growth medium containing 200 µM uniformly U-^13^C labeled palmitate (Cambridge Isotope Laboratories, CLM-409-0.5) and 5% delipidated FBS (Gemini Bio-products, 900-123). For the glucose and glutamine tracing experiments, the transfected cells were incubated for 72 h with DMEM/F-12 growth medium containing either 12mM uniformly U-^13^C labeled glucose (Cambridge Isotope Laboratories, CLM-481-0.5) or 2mM uniformly U-^13^C labeled glutamine (Cambridge Isotope Laboratories, CLM-1822-H-0.5), respectively. The sample preparation and processing were performed as described in the untargeted metabolomic section above. No internal standards were added into the isotope tracer samples. UHPLC-HRMS and metabolomics data normalization were performed as described in the untargeted metabolomic section above. Data analysis was performed using El-MAVEN.

### In vitro RNA pull-down coupled with protein detection

For *in vitro* RNA pull-down, the magnetic RNA-protein pull-down kit (Pierce, 20164) was used according to the manufacturer’s instructions. The *in vitro* transcribed lncRNA and its antisense strand were end-labeled with desthiobiotin, and 50pmol of biotin-labeled RNAs were incubated with 50 µL streptavidin magnetic beads for 30 min at 25 °C with agitation. Streptavidin magnetic bead-bound RNAs were incubated with CA1D cell lysate (50 to 300 µg) overnight at 4°C with gentle end-to-end rotation. Beads were then washed with 1X Wash Buffer and resuspended in 50 µL of Elution Buffer provided in the kit. The eluted RNA-bound proteins were separated by SDS-PAGE and detected with anti-CPT1A monoclonal antibody (Abcam, ab128568, 1:1000).

### Detection of carnitine levels via targeted metabolomics

#### Sample preparation

CA1D cells were transfected with either siControl or siTROLL-8. 48 h post-transfection, cells were harvested and washed 2-3 times with ice-cold PBS. Samples were spiked heavily with labeled internal standards (10 labeled acyl-carnitines). Cold 80% methanol extraction solvent was added to the sample for protein precipitation. After vortexing, the samples were incubated at -80 °C, followed by centrifugation. The supernatant was transferred to new Eppendorf tubes and dried. The dried protein pellets were measured by Bradford and then re-dissolved in 10 µL of 80% methanol for analysis.

#### Reagents and chemicals

Ammonium hydroxide, ammonium carbonate and all the 12 acyl-carnitines (L-carnitine, O-acetyl-L-carnitine, butyryl-L-carnitine, hexanoyl-L-carnitine, octanoyl-L-carnitine, decanoyl-L-carnitine, lauroyl-L-carnitine, myristoyl-L-carnitine, palmitoyl-L-carnitine, stearoyl-L-carnitine, oleoyl-L-carnitine, hydroxybutyryl-L-carnitine) were obtained from Sigma Aldrich and used for method development. LC-MS grade water, methanol and acetonitrile were from VWR. L-carnitine (trimethyl-D9, 98%), acetyl-L-carnitine (N-methyl-D3, 98%), butyrl-L-carnitine (N-methyl-D3, 98%), hexanoyl-L-carnitine (N-methyl-D3, 98%), octanoyl-L-carnitine (N-methyl-D3, 98%), decanoyl-L-carnitine (N-methyl-D3, 98%), lauroyl-L-carnitine (N,N,N-Trimethyl-D9, 98%), myristoyl-L-carnitine (N,N,N-Trimethyl-D9, 98%), palmitoyl-L-carnitine (N-methyl-D3, 98%) and stearoyl-L-carnitine (N-methyl-D3, 98%) were purchased from Cambridge Isotope Lab and used for metabolite quantification.

#### LC-MS metabolomics

UHPLC-MS was performed using a Vanquish LC (Thermo Fisher Scientific) interfaced with a Q Exactive FOCUS mass spectrometer (Thermo Fisher Scientific). Chromatographic separation was performed on an ACQUITY UPLC BEH Amide column (2.1 mm x 150 mm, 1.7 µm particle size, Waters). The mobile phase A was 10 mM ammonium carbonate and 0.05% ammonium hydroxide in water, and the mobile phase B was 100% acetonitrile (total run time of 15 min). The column temperature was set to 30 °C, and the injection volume was 2 µL. The Single Ion Morning (SIM) is performed in positive mode. Skyline was used for data analysis.

### Cell-fractionation and mitochondrial isolation

2 x 10^7^ CA1D cells were used for the extraction of cytoplasmic and mitochondrial proteins using the Mitochondrial Isolation Kit for Cultured Cells (Thermo Fisher Scientific, 89874) according to the manufacturer’s instructions. In all, 50ug to 300ug of the cytoplasmic and mitochondrial fractions were analyzed by SDS-PAGE, and the following primary antibodies were used: anti-CPT1A (Abcam, ab128568, 1:1000), anti-alpha-tubulin (Abcam, ab52866, 1:1000), and anti-VDAC (Abcam, ab154856, 1:1000).

### Identification of CPT1A-interacting proteins via liquid chromatography-mass spectrometry (LC-MS/MS)

1.5 x 10^6^ CA1D cells were transfected with either siControl or siTROLL-8. 48 h after the transfection, the cells were lysed in IP lysis buffer (Pierce, 87787) and the IP assay was performed using 4 µg of either CPT1A antibody (Proteintech, 15184-I-AP) or normal rabbit IgG (Santa Cruz, sc-2027) as a negative control. Immunoprecipitated proteins were extracted, digested with trypsin, and analyzed via LC-MS/MS^2^, and the identified peptides are listed in Supplementary Table 4.

### Co-immunoprecipitation (CoIP) assay

CA1D cells were transfected with either siControl or siTROLL-8 and collected 48 h post-transfection. The cells were then lysed, and the CoIP assay was performed in IP Lysis Buffer (Pierce, 87787). 2 µg of each of the following primary antibodies was utilized per sample: CPT1A (Proteintech, 15184-1AP) and normal rabbit IgG (Santa Cruz, sc-2027) as a negative control. The interaction was then detected via western blot using the following primary antibodies: ACSL1 (Invitrogen, FEMA550618, 1:1000), CPT1A (Abcam, ab128568, 1:1000), and VDAC1 (Santa Cruz, sc-390996, 1:100).

### Soft agar colony assay

CA1D were transfected with siControl, siACSL1, siCPT1A, siTROLL-8, and siVDAC1. After 48 h, the cells were trypsinized and resuspended in a top layer of their culture medium with 0.3% agarose (Thermo Fisher Scientific, BP160) at 1 × 10^5^ cells per well in triplicate in six-well plates and plated on a bottom layer of culture medium containing 1% agarose. 1 mL of medium was added to the top layer of each well and changed every 3 days. After 5 weeks the colonies were counted with a 20X objective on a Zeiss Observer.Z1 microscope.

### Kaplan-Meier curves of TCGA data

To assess the clinical significance of CPT1A and VDAC1 in breast cancer, we first downloaded the TCGA breast cancer isoform expression dataset transcriptome profiles^22^ from the Broad Institute and clinical dataset transcriptome profiles from cBioPortal. All analyses were divided into two cohorts according to the median expression of their respective RNA (high vs. low expression). The expression class was defined as follows: expression value = 0 was excluded, 0 < expression value ≤ median was defined as low expression, expression value > median was defined as high expression.

### Statistical analysis and reproducibility

All the experiments are representative of at least three independent replicates. Data collection was performed with Microsoft Excel, and data analysis was performed using GraphPad Prism, Oncotopix® (Visiopharm), and R software. Gels, blots, and images are representative of three independent experiments giving similar results.

